# Testing competing models of dorsal anterior cingulate

**DOI:** 10.1101/273706

**Authors:** Eliana Vassena, James Deraeve, William H. Alexander

## Abstract

Recent theories have attempted to provide unifying accounts of dorsal anterior cingulate cortex (dACC), a region routinely observed in studies of cognitive control and decision-making. Despite the proliferation of frameworks, rigorous empirical testing has lagged behind theory. Here we test competing predictions of three accounts of dACC using a simple value-based decision-making task. We find that the Predicted Response-Outcome model provides an integrative and parsimonious account of our results. Our results highlight the need for increased emphasis on empirical tests of theoretical frameworks.

---

Activity in dorsal anterior cingulate cortex (dACC) and surrounding regions in medial prefrontal cortex (mPFC), is routinely observed in neuroimaging studies of cognitive control and decision-making^1^. Consequently, a number of computational accounts have been developed in the last two decades to describe the role and function of dACC^2–6^. Formal tests between competing models are critical in order to drive efficient acquisition of knowledge, and recent efforts in this direction^7,8^ have done much in advancing the debate regarding brain function.

In this manuscript, we focus on the predictions of three models of dACC. The Choice Difficulty (CD) account^9^ states that dACC activity codes for choice difficulty (deriving from value similarity between available options), and has been advanced as a broadly-applicable model for explaining effects observed in dACC, including, e.g., as an alternative to the proposal that dACC calculates the relative value of foraging^10^. A second view, the Expected Value of Control (EVC) model^11^ states that dACC integrates value and cost information in order to derive an optimal control signal (balancing task demands against prospective rewards). Finally, the Predicted Response-Outcome (PRO) model^12^ states that dACC learns to predict the likely outcomes of actions and signals deviations between expected and observed outcomes. Simulations of these models in the context of value-based decision-making predict substantially different patterns of dACC activity (Figure 1A; see Online Methods). Under the CD model, dACC should track the similarity of options: activity should be maximal for options with similar values, and decrease as the value difference increases. Predictions of the EVC model were derived in two ways. Using only the equations presented in Shenhav et al 2013, the EVC model (EVC1) predicts a pattern opposite to that of the CD model: dACC activity should be minimal when options are similar (exerting control is not worth it as both option are equally valuable), and increase with increasing value difference between options (reflecting increased value for exerting control as the value of one option increases). If we include the idea (described but not formalized in Shenhav et al 2013) that, as decisions become easier due to large value differences, the value of additional control decreases, the EVC model (EVC2) predicts an ‘M’ shaped pattern. The optimal control signal for similar options is low (as in EVC1, additional control does not increase the value of outcomes), as is the control signal for extremely different values (no control is required to make a choice). Finally, the PRO model predicts a ‘W-shaped’ pattern in dACC activity (Figure 1), derived from the negative component of prediction error (PE), i.e. negative surprise^12^. PEs are commonly observed throughout the brain when actual events differ from expected or intended events. For options with similar values, evidence for choosing either option is approximately equal (i.e., a person may equally intend to select either option) and therefore either choice entails surprise related to the unmet partial intention to select the alternate option. Surprise signals also apply at the onset of trials when options are revealed: on trials with extreme differences in the value between options, surprise derives from the unexpected deviation of option values from their long-run average. The ‘W’ pattern predicted by the PRO model thus derives from standard PE calculations at the presentation of options and generation of a response (cf. Fig. S3, Online Methods).

**Figure 1.**
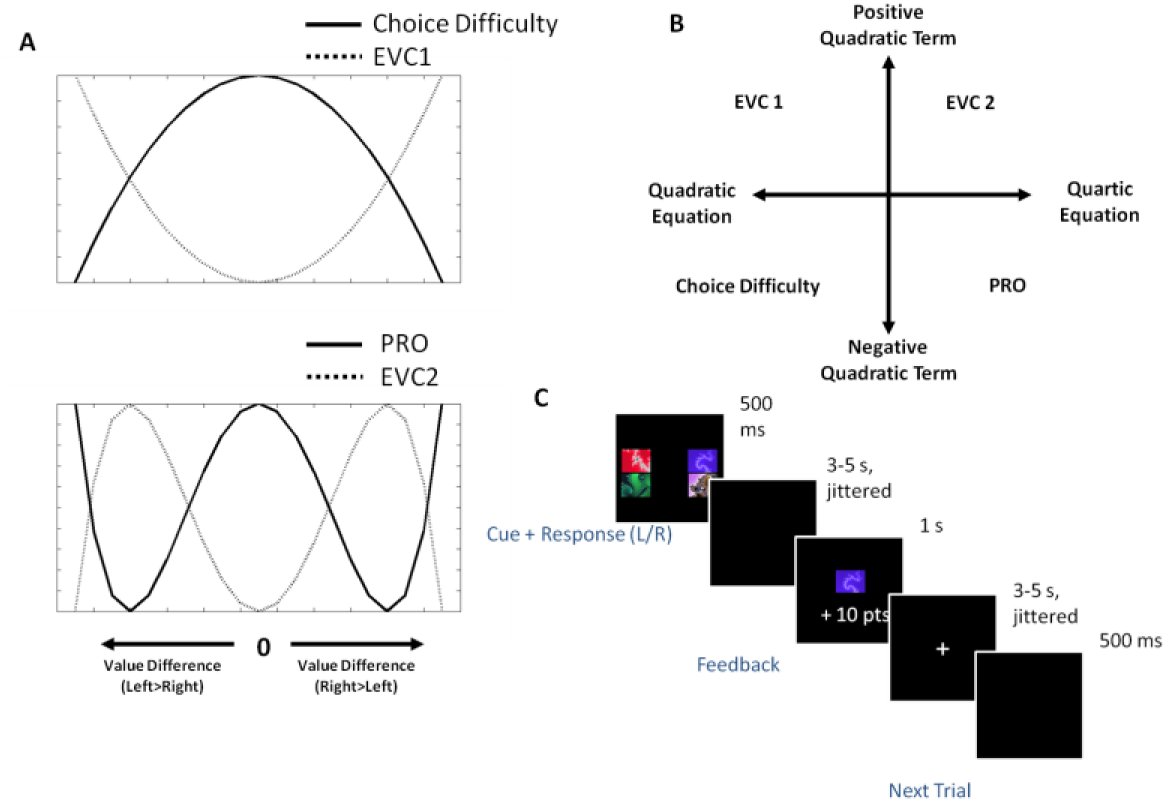
A) Qualitative predictions of the models for dACC activity. Choice Difficulty and EVC1, predict a quadratic curve. The PRO predicts a W-shaped curve. EVC2 predicts an M-shaped curve. B) Summary of analysis rationale. Each model makes predictions that can be discriminated along two dimensions. C) The speeded decision-making task. Subjects choose between two options (left/right), with a 50% chance of receiving the points associated with each of the images for the selected option.

A convenient way to summarize these competing predictions is as polynomial equations (Fig. 1A), which transcend model-specific implementation details. The predictions of both CD and EVC1 can be characterized as quadratic polynomials^9^, the difference between the two being the sign on the coefficient for the quadratic term (U-shaped for EVC1, inverted U for CD). The predictions of the PRO model and EVC2 are best characterized by a quartic (fourth-order) polynomial, also with opposite signs on the coefficient of the quadratic term (negative for the PRO model, positive for EVC2).

We can thus distinguish amongst the model predictions as follows (Fig 1B): first, comparisons of Akaike’s Information Criterion (AIC) values for fits of polynomial equations to neural data provide an estimate of whether the data are better explained by quadratic (CD/EVC1) or quartic (PRO/EVC2) curves. Second, the sign on the quadratic term of the best-fit polynomial equation distinguishes between remaining models (negative (CD/PRO) or positive (EVC1/EVC2, Fig 1B).

Model predictions were tested by having human subjects perform a speeded value-based decision task while undergoing fMRI (fig 1C). Data were modeled using a general linear modeling approach (GLM), and 2 GLMs were constructed for our analyses (See Online Methods). In GLM1, 11 regressors modeled the binned differences in the expected value of the two options (left/right). In GLM2, 8 regressors modeled left/right RT bins with an equal number of trials per bin. For each voxel in an anatomically-defined ROI including mPFC (Fig 2A), dACC, and SMA, polynomial equations (1st through 4th order) were fit to the estimated beta values for each voxel across all subjects for each regressor in GLM1 and GLM2.

**Figure 2.**
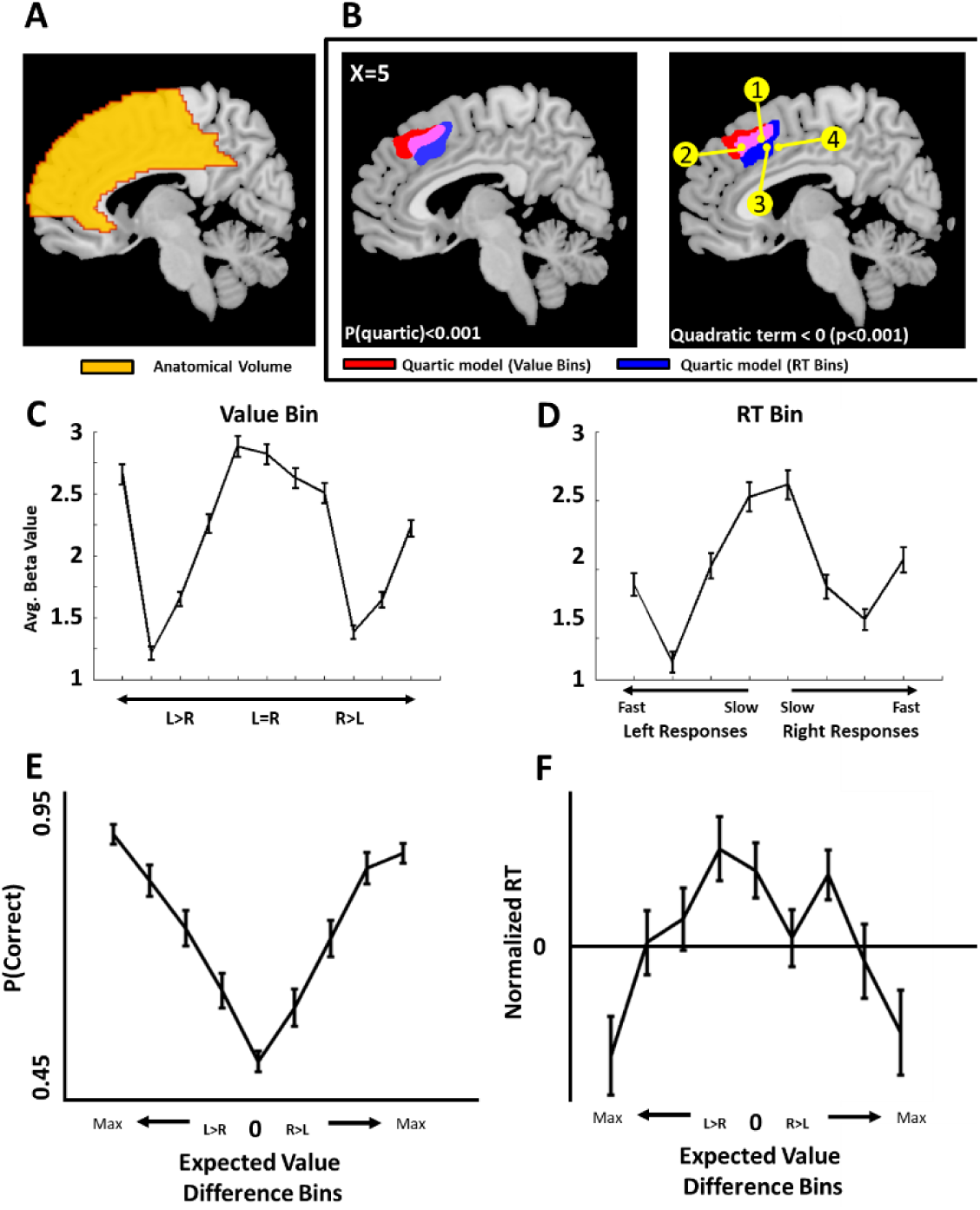
A) Anatomically defined region including dACC, MPFC and SMA where model predictions were tested. B) Only signals associated with quartic polynomial equations (left) were identified in the region, consistent only with the PRO and EVC2 models. The best-fit equation for voxels in this region included a quadratic term that was significantly less than 0 across subjects (right), consistent with the PRO model, and inconsistent with the EVC2 model. Yellow dots indicate peak voxels reported in previous studies observing dACC activity related to Choice Difficulty (1 & 2) 9,14 and Value Difference (3&4) 10 C) Average beta weights for voxels passing a threshold of p<0.001 within the anatomical region binned by value and D) reaction time. E) Subjects’ ability to correctly select the option with the higher expected value improved as a function of the difference in expected value between options, and F) reaction times decreased as a function of value difference. Behavioral results therefore rule out a possible explanation for increased dACC activity for extreme trials as being due either to increased error rate or increased reaction time.

To assess which polynomial equation better explained the data, Akaike weights^13^ were computed for the AIC value obtained in order to derive a probability for that each equation amongst those considered best explained the data. The probability for each equation at each voxel was thresholded at an Akaike weight of > 0.999 (equivalent to p=0.001). To calculate statistics with volume-based corrections, the best-fit polynomial equations for each voxel passing threshold were averaged and regressed against the BOLD signal. For both value-binned (GLM1) and RT-binned (GLM2) values, a cluster of voxels surviving cluster correction (voxel threshold of p=0.001, cluster-level FWE = 0.05) was observed in dACC/mPFC for the quartic polynomial (Fig. 2B, Left). No effects consistent with other polynomial equations (linear, quadratic, cubic) were observed in the region, even at a lenient threshold (p=0.05, uncorrected). Thus, activity in dACC/mPFC is best explained by a quartic polynomial, consistent with the PRO and EVC2 models.

To distinguish between PRO and EVC2 models, the sign on the quadratic term for the voxels identified in our first step was tested; EVC2 predicts a positive quadratic term, while the PRO predicts a negative quadratic term. Only voxels with a quadratic term significantly less than 0 (p<0.001) were identified, consistent with the PRO model and inconsistent with EVC2 (Fig. 2B, Right). In summary, these results favor the PRO model over the EVC and CD models in explaining dACC activity during value-based decision-making with time pressure.

Additional support for the PRO model in our data comes from analysis of activity at the time of feedback. As described above, under the PRO model dACC activity is related to the “suprisingness” of a salient event. In our previous analyses, we limited ourselves to the interval starting from the onset of a trial to the generation of a response, primarily because the CD and EVC models do not make clear predictions regarding dACC activity absent the requirement for generating behavior. If our data are consistent with the PRO model account, dACC activity following feedback should correlate with the (unsigned) feedback PE. Regression of the per-trial PE for each subject on BOLD data yields a significant cluster (Fig. 3; peak voxel −4, 38, 42, cluster-corrected p(FWE)<0.001, voxel extent = 2029, voxel-wise threshold = 0.001) in dACC. The PRO model thus accurately accounts for dACC activity over the course of the entire trial in our speeded decision-making task.

**Figure 3.**
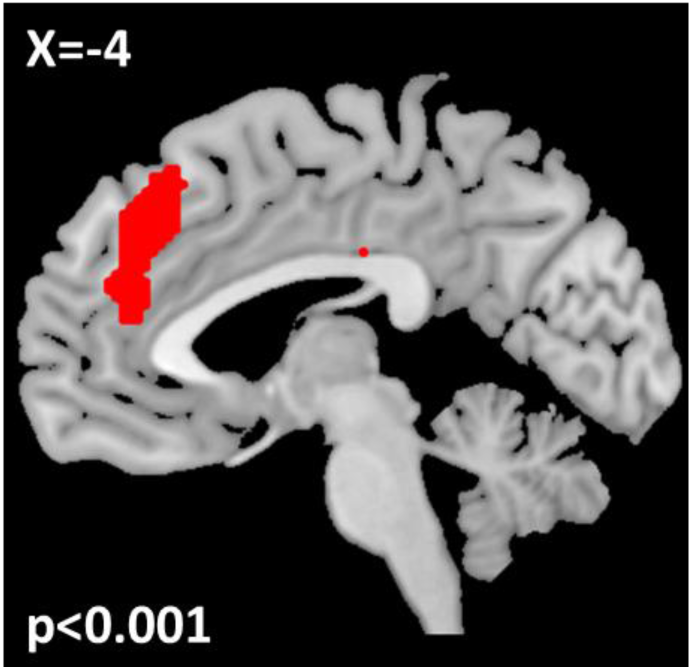
DACC activity correlates with unsigned prediction errors following task feedback consistent with the PRO account

The PRO model accounts for quartic effects observed in dACC through the single mechanism of *negative surprise,* derived from standard calculations of prediction errors, and provides a strong prediction regarding the time course of dACC activity producing the ‘W’ pattern (cf. Fig. S3, Online Methods). At stimulus presentation and prior to response, the PRO model predicts increased activity as a function of value difference. Following a response, it predicts increased activity as a function of value similarity. It may be noted that the pattern of activity observed in our data might be captured by a combination of signals generated by the EVC and CD models; however, it is not clear what additional explanatory power would be provided by such a dual-mechanism account, nor under what circumstances such an account could be falsified. The PRO model, in contrast, provides a parsimonious and testable explanation for the pattern observed in our data. Future work should therefore focus on elucidating the temporal dynamics of dACC activity using methods with higher temporal resolution in order to test these specific predictions. More generally, direct empirical investigation is essential for adjudicating amongst competing accounts of brain function and developing a more complete understanding of the neurobiological mechanisms underlying cognition; the current work adds to the growing emphasis on developing and testing models of the brain ^7–9^.

## Acknowledgments

We thank Clay Holroyd, Joshua Brown, Tom Verguts, and Mathias Pessiglione for useful discussion. WHA was supported by FWO-Flanders Odysseus II Award #G.OC44.13N.EV was supported by the Marie Sklodowska-Curie action with a standard IF-EF fellowship, within the H2020 framework (H2020-MSCA-IF2015, Grant number 705630).

## Online Methods

### Participants

Twenty-three healthy volunteers participated in this experiment (10 males), with a mean age of 23 ±2, ranging between 19 and 28. The Ethical Committee of the Ghent University Hospital approved the experimental protocol. All participants signed an informed-consent form before participating in the experiment, and filled in a safety checklist to exclude contraindications for participations, and neurological or psychiatric conditions.

### Experimental procedure

Upon arrival, participants filled in and signed the informed consent and the pre-scanning checklist. Subsequently, they underwent a training session outside the scanner. The training consisted of performing a series of decisions. On each trial, participants were presented with two fractal images on the screen, one on the left and one on the right (1000ms). Participants could select one image by pressing the left or right response key on the keyboard. After 500ms, feedback was given, showing the selected image and the amount of points obtained in that trial (1000ms). After a blank screen (1000ms) the following trial started. There were 8 images in total. Each image was associated with a fixed amount of points, ranging (in 10 point increments) between 10 and 80 points. Participants were instructed to learn the amount of points associated with every image. They were informed that the task during the subsequent scanning session would be different, but the same images would appear, and each image would be associated with the same amount of points as in the training. They were therefore specifically encouraged to remember this association. Participants were also informed that all points gathered during the scanning session would be converted into money, which they would be paid at the end of the experiment in addition to the amount paid for participation. At the end of the training session all participants confirmed that they learned the amount of points associated with each image, and that some images delivered more points than other images. Importantly, the amount of points associated with each fractal image was changed across participants according to 8 possible randomizations in order to control for visual features of the images.

### Speeded value-based decision-making task

Participants performed a speeded value-based decision-making task while undergoing fMRI. The task started with a short training session, to acquaint participants with the procedure and the response buttons in the scanner. A total of 20 training trials were performed. On each trial, participants were presented with 4 fractal images, two on the left side of the screen, and two on the right side of the screen (500ms; Figure 1C). They were instructed to select options (left or right side of the screen) in order to maximize the number of points earned. Of the chosen side, they had equal probability of receiving one of the two presented images as feedback, and the corresponding amount of points, as feedback. Following a response, a blank screen was presented for a jittered variable interval (randomly selected, range 3000-5000ms, mean 4000ms). A feedback display followed, showing the image selected by the computer among the images on the chosen side, with the corresponding amount of points obtained by the participant (500ms). Feedback was followed by a randomly jittered interval, ranging between 3000 and 5000ms (mean 4000ms). After completing the training, participants were asked if the points associated with each image corresponded with what they had learned in the training outside the scanner, and every participant so confirmed. Subsequently, the task started, with same timing and procedure. The task consisted of 160 trials in total in one block, lasting approximately 25 minutes.

Activity in dACC is known to correlate with uncertainty and choice variance^1^. The model-derived predictions regarding dACC activity (see below) suggest that, under certain circumstances, dACC activity may increase as the difference between options increases. In the speeded decision-making task, the possible outcomes for each option are selected at random from the set of 8 outcomes as described above. As a consequence, the variance within each option decreases as the variance between options increases (Fig. S1), and we can rule out the possibility that increase in variance drives increases in dACC activity for extreme trials.

**Figure S1.**
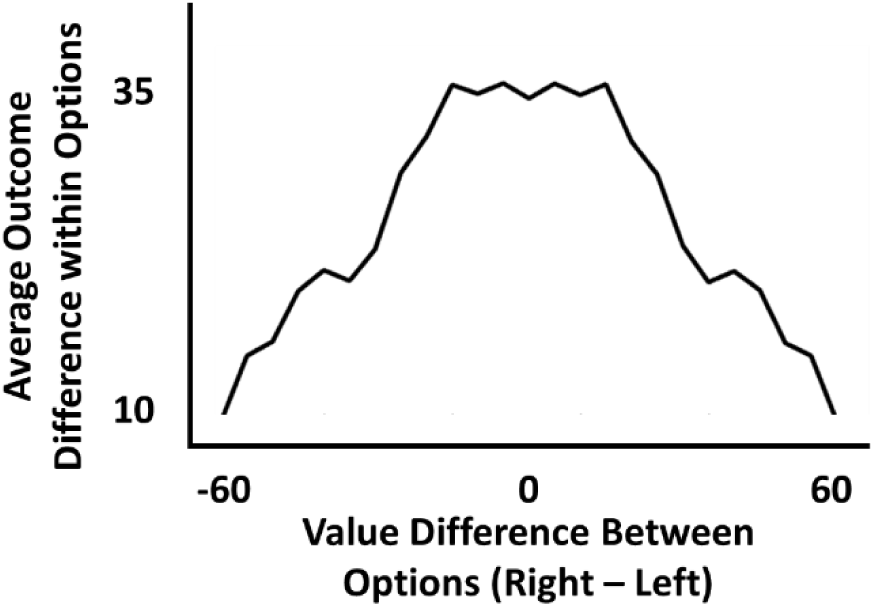
Within-option value difference as a function of between-option value difference

### fMRI data acquisition

Data were acquired using a 3T Magnetom Trio MRI scanner (Siemens), with a 32-channel radio-frequency head coil. In an initial scanning sequence, a structural T_1_ weighted MPRAGE sequence was collected (176 high-resolution slices, TR = 1550 ms, TE = 2.39, slice thickness = 0.9 mm, voxel size = 0.9 × 0.9 × 0.9 mm, FoV = 220 mm, flip angle = 9°). As second sequence, functional images were acquired using a T_2_* weighted EPI sequence (33 slices per volume, TR = 2000 ms, TE = 30 ms, no inter-slice gap, voxel size = 3 × 3 × 3mm, FoV = 192 mm, flip angle = 80°). On average 760 volumes per participants per task were collected. Each task lasted approximately 25 minutes.

### 2.5 fMRI data analysis

The first 4 volumes of each functional run were discarded to allow for steady-state magnetization. The data were preprocessed with SPM 8 (http://www.fil.ion.ucl.ac.uk/spm). Images were realigned to the first image of the run. The structural T_1_ image was coregistered to the functional mean image for normalization purposes. Normalization was performed through the unified segmentation and nonlinear warping approach implemented in SPM8. Functional images normalized to the MNI template (Montreal Neurological Institute). Resulting functional images were smoothed with a Gaussian kernel of 8 mm full width half maximum (FWHM).

### Speeded value-based decision-making task

For each single subject, a General Linear Model (GLM) approach was applied in order to identify condition-specific activation. In a first GLM (GLM1), trials were divided in 11 different bins as a function of the value difference between the two sides of the screen (5 bins for the left side average value > right side average value ranging between 55 and 10 points difference, 1 bin for trials where both sides were close in value (−5 to +5 average point difference for Right-Left options), 5 bins for the right side average value > left side average value, ranging between 55 and 10 points difference). For each regressor, a parametric modulator with RT at the current trial was added. An additional regressor was added to model responses over time limit (misses). Twelve regressors were added to model feedback in each condition and in misses. Six more regressors were added to account for motion (X,Y, Z translation, pitch, yaw, and roll). In a second GLM (GLM2), trials were divided in 8 different bins as a function of RT and side. The purpose of this binning procedure was to equalize the number of trials per bin, and to ensure that possible increases in activity for extreme value difference could not be attributed to increased RT. For each regressor the RT on current trial was added as a parametric modulator. One regressor was added for responses over time limit. Nine regressors were added to model feedback in each condition and feedback for misses. Six regressors were added to account for motion as above.

Because each computational model under consideration predicts effects specifically related to dACC and medial prefrontal cortex (mPFC), we restricted our analyses to an anatomically-defined region including dACC, supplementary motor area (SMA), and superior mPFC, defined anatomically using the wfupickatlas toolbox (dilation=2). The predictions of each model considered (see below) can be described in a straightforward fashion by polynomial equations. Therefore, for all voxels within this region, beta values estimated for that voxel at the 1st level, and for all subjects, for each regressor in GLM1 were used to fit polynomial equations (linear, quadratic, cubic, and quartic). Following polynomial fits, an AIC value was computed for each polynomial equation, and AIC values were used to compute Akaike Weights^2^ yielding a value between 0 and 1 for each model indicating the probability that model is the best of all models under consideration. For each voxel, therefore, a value was obtained indicating the probability that the beta values at that voxel were best explained by each of the 4 polynomial equations. These values were thresholded at 0.999 (equivalent to p<0.001), and for all voxels surviving this threshold, the best-fit polynomial equation was determined, averaged over all surviving voxels, and regressed against the BOLD signal to compute whole-brain statistics. These steps were repeated for GLM2 (binned by RT).

To analyze feedback-related activity, a third GLM (GLM3) was created in which one regressor, corresponding to the onset of task feedback was modeled on each trial. A parametric modulator reflecting the (objective) unsigned prediction error (calculated from global task contingencies and based on subjects’ trial-by-trial choices) was included for the feedback regressors. A second regressor was used to model non-response trials, and 6 motion regressors were included as in GLMs 1 & 2.

## Modeling Methods

All scripts used to derive predictions from the various computational models are available online at https://github.com/modelbrains/EVC-Simulations

### Expected Value of Control (EVC1)

Equations for the Expected Value of Control^3^ (EVC) model were taken from Shenhav et al. (2013). Specifically, for 1-step choice tasks, the EVC model is specified by the equations:

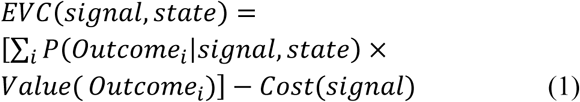

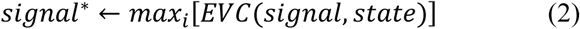

Although many aspects of the EVC model, such as the form of cost functions related to effort and the relationship between effort and probability, are unspecified, there are three clear commitments the model makes. First, increased control is positively and monotonically related to the cost of control: exerting more effort to make a choice *always entails* additional costs. Second, the level of control is positively and monotonically related to the probability of success: exerting more effort to make a choice *always increases* the probability of successfully making that choice. Finally, the EVC model states that dACC activity is proportional to the intensity of the optimal control signal (eq. 2).

In order to derive predictions from the EVC model using only the equations provided in Shenhav et al., 2013 (EVC1 in the main text), we simulated the EVC1 model on 100,000 trials of the speeded decision task. Since the EVC model leaves the specification of the functions (cost function and p(success|effort)) noted above ambiguous, the model was simulated multiple times using different functional forms that nonetheless satisfy the central commitments described above. Candidate signal costs (eq. 1) were modeled in the range [0 5] for each of the possible actions in the speeded decision-making task. The probability of the model selecting an option given a certain effort level was modeled as a sigmoid function

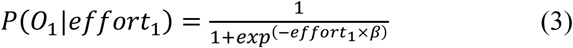

where *effort*_*1*_ is the level of effort exerted in order to realize outcome *O*_*1*_ and β is a scaling parameter that determines how quickly the sigmoid function asymptotes for increasing effort levels. Because there were only two possible options to choose from, P(O_2_|effort_2_) was modeled as 1-P(O_1_|effort_1_). To accommodate possible nonlinearities in the cost function, the model was simulated with cost functions exponentiated by values of 0.5, 1, and 2. As with the cost function, the probability function was simulated in three different ways to account for possible non-linear relationships between effort and probability. Specifically, the parameter β was assigned values of 0.1, .5 and 10. Crossing the 3 cost function manipulations with the 3 probability function manipulations yields 9 simulated conditions. Regardless of the particular form of cost and probability functions, however, the qualitative predictions of the EVC1 model are identical in that they all produce a U-shaped curve (cf. figure 1A, main text) centered at a value difference of 0. The predictions of the EVC1 model are therefore approximated by a quadratic polynomial with a positive sign on the quadratic term.

### Expected Value of Control 2 (EVC2)

An additional aspect of the EVC model, not specifically formalized in Shenhav et al., (2013) is that, as decisions become easier, control costs decrease. In the speeded decision task, options with large value differences may therefore represent easy decisions in which control is not necessary since the response is largely driven by bottom-up processes. In order to derive predictions from the EVC model incorporating this intuition, we altered eq. 3 to include a shifting set point for determining choice probabilities

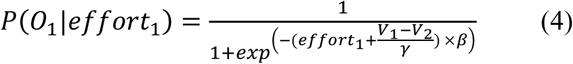

where *V*_*1*_ and *V*_*2*_ are the expected value of options 1 and 2, respectively, and γ scales the value difference between the 2 options (β *= 1*). The effect of this addition to the model for choice probability is to change the effort point at which the model is indifferent between options. When *V*_*1*_ *= V*_*2*_, eq. 4 is the same as eq. 3. As *V*_*1*_ becomes greater than *V*_*2*_, the point at which the model is equally likely to select either option shifts toward *V*_*2*_ - the model would need to exert effort favoring the selection of *V*_*2*_ in order to have an equal probability of selecting either option. Put another way, it is more likely that the model selects *V*_*1*_ with no effort expended, and this probability increases with the difference in value between options. Thus eq. 4 captures the intuition that choice becomes easier as the difference in values increases.

**Figure S2.**
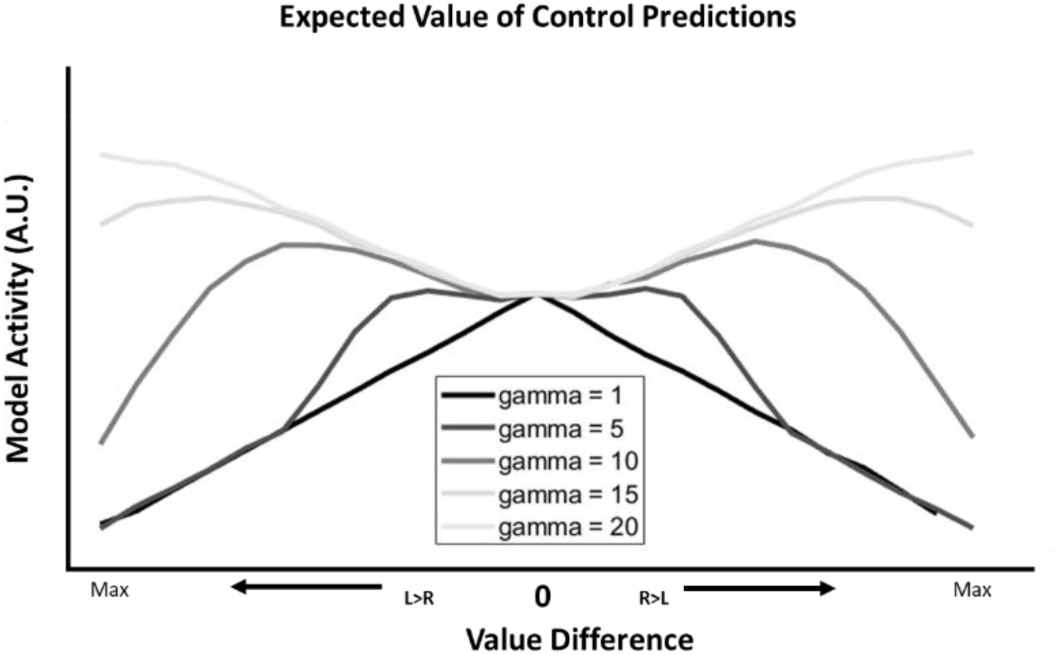
Predictions for the EVC2 model as a function of temperature (gamma)

The predictions derived from EVC2 are slightly more nuanced than those for EVC1. For very low values of γ, the influence of value differences overrides the effect of effort - any value difference becomes the dominant factor in determining choice. Here, the predictions of EVC2 correspond to those of the choice difficulty model (see below), with the greatest predicted activity for choices close together, and the least activity for options with large value differences. Conversely, when γ is very high, predictions of EVC2 correspond to those of EVC1 - here, effort is the decisive factor in choice behavior, and the rationale derived from simulations of EVC1 applies. Of interest, however, is the behavior of the model for intermediate values of γ. As γ increases from low to high values, the model predictions assume an ‘M’ shape. Notably, at no point in this progression does the predicted activity assume a ‘W’ shape. Thus the EVC2 model may be consistent with either EVC1 or CD, but not with the PRO model (see below).

The ‘M’ shape has minima at the following points: when the value difference between options is most extreme, and when the options are approximately equally valuable. Intuitively, this arises from the sensitivity of the optimal control signal both to changes in incentives as well as the costs required to realize the incentives. At extreme value differences, incentives are high, associated with a stronger control signal. However, the decision for extreme value difference trials is relatively easy, meaning that the costs of control are low, requiring a weaker control signal. Conversely, when both options are equally valued, the incentive for choosing one over the other is minimal, implying a weak control signal, while the decision costs are higher due to the similarity of the options, implying a strong control signal. The minima in the ‘M’ pattern therefore correspond to situations in which the control signal intensity depends entirely only on one factor in the decision process, i.e., only costs or reward. The peaks of the ‘M’, in contrast, reflect those trials in which both cost and reward substantially contribute to the reward signal.

### Applicability of EVC to speeded decision-making and derivation of predictions

As indicated above, several aspects of the EVC model are left un- or under-specified. Nevertheless, previous statements in publications describing the EVC model informed our choice of a speeded value-based decision-making task to test predictions of the EVC model. Here, we specifically cite those statements justifying our choice of task, as well as the manner in which we derive predictions from the model. All **emphasis** is not in the original, but included to highlight points relevant to the current manuscript.

**1) EVC applies to economic tasks with control requirements.** In Shenhav et al., 2014, the authors state:

> “We recently described an integrative theory of dACC function, which proposed that the dACC is responsible for estimating the expected value of control-demanding behaviors (EVC) and selecting which to execute. Like the KBMR theory, **this theory predicts that dACC activity should track the expected reward** for engaging in non-default behavior**, inasmuch as this can be considered to be control-demanding**.” ^4^

**2) Time pressure is a control-demanding experimental manipulation.** In Shenhav et al., 2013, the authors state:

> “The EVC model proposes that dACC mediates these adjustments, by monitoring for the conditions that require them, and specifying the necessary adjustments for other systems responsible for implementing them. This makes two predictions: first, **that dACC should be responsive to conditions indicating the need to adjust control intensity;** and, second, that it should be associated with the engagement of neural systems responsible for implementing these adjustments (i.e., the regulative function of control).
>
> “There is extensive evidence in support of the first prediction, indicating that **dACC is responsive to conditions** requiring adjustments of threshold and/or response bias, **such as increases in time pressure** and changes in prior probabilities…” ^3^

**3) DACC activity and the optimal control signal.** The authors of EVC indicate a direct relationship between the optimal control signal and dACC activity. Specifically, in Shenhav et al., 2013, they write:

> “Under plausible assumptions about the shape of the payoff and cost functions (see Kool and Botvinick, 2012**), the optimal control signal intensity will rise with the magnitude of task incentives** (see Figures 4A and 4B). **This predicts that dACC activity should grow both with task difficulty and with the stakes associated with task performance.**” ^3^

The speeded decision-making task used in this study involves making economic choices within a relatively short time window, a context in which the authors of EVC specifically state that dACC activity should track reward **(1)** since it involves making responses under time pressure **(2)**, a control-demanding manipulation specifically noted in Shenhav et al., 2013. Our implementation of the EVC model derives predictions of dACC activity based on the intensity of the optimal control signal **(3)**; the positive relationship between the optimal control signal intensity and dACC activity is specifically endorsed by the authors of EVC.

Finally, in informal discussions relating to our simulations of the EVC model, it was questioned why a model with perfect information regarding available choices, as is the case in our initial simulations, should require control in the first place. In response, we note that our simulations were conducted specifically with the speeded decision-making task in mind, and, as has been noted, time pressure is specifically identified as a control-demanding manipulation. That is, even if perfect information regarding options is available, control is still necessitated in order to govern an ongoing response process that, due to time pressure, needs to be initiated and completed relatively quickly after stimulus presentation. Additionally, even though our initial simulations assumed perfect information, this is not required in order to derive the predictions we attribute to the EVC model. In additional simulations of the EVC model, we introduce estimation noise as follows: on each trial, rather than calculating the veridical expected value of each option, we instead took the average of 5 random samples drawn from a uniform distribution bounded by the minimum and maximum possible value for each option. The results of these simulations reproduce the qualitative patterns observed for simulations of the EVC2 model above.

### Choice Difficulty

The Choice Difficulty model^4^ is formulated as the negative absolute value of the difference in subjective value between two options:

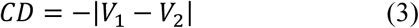

Subjective value may be biased due to task demands, such as the presence of a default response that biases subjects against selecting a non-default option. Additionally, it is possible that additional terms may further inform the calculation of value difference - e.g., exponential terms that add a nonlinear component to value. Nevertheless, the CD model is quite clear that dACC activity should be maximal for subjective value differences near zero, and minimal for extreme value differences. That is, regardless of response bias, the CD model can be approximated by a quadratic polynomial equation with a negative sign on the quadratic term.

### Predicted Response-Outcome Model

The Predicted Response-Outcome^5^ (PRO) model, in contrast to the previous models, does not interpret dACC function as specifically learning about value information, but rather about the likely outcomes of actions. Nevertheless, in our initial publication of the model^5^ as well as in applications of the model to substance dependence^6^, we noted that value could influence the salience of an observed outcome - high value outcomes are more salient than low value outcomes, and thus the predictions learned by the PRO model could reflect a biased estimate of the likelihood of outcomes depending on subjective value. We simulated the PRO model on the speeded decision-making task using our previously published parameter set. Inputs to the model were the 8 possible task stimuli, and two outcomes were modeled for each option, with the value of the outcome modeled as the total number of points earned by selecting an option. Following feedback, model activity was allowed to decay back to baseline prior to the onset of the next trial. Activity (negative surprise) was recorded from the model beginning at the presentation of the two options in the task until 20 iterations (200ms) following the generation of a response. The model was simulated on 10 different experimental runs of 3000 trials each, and data were recorded after the first 500 trials to allow the model time to learn task contingencies. Predictions for the PRO model are ‘W’-shaped, with predicted activity in dACC high for conditions in which the value of 2 options is similar, as well as conditions with extreme value differences.

The PRO model interprets dACC activity as being primarily related to the calculation of prediction error (specifically, the negative component of a prediction error), a quantity that has frequently been applied to the interpretation of neural data. In the speeded decision task, prediction errors (PEs) occur at three points in each trial. First, over the entire experiment, the mean value of both the left and right options is approximately 45 points – that is, at the onset of a trial, and prior to seeing the fractal images, it can be predicted that, on average, each option will be worth around 45 points. When the fractal images are presented, the actual value of each option can be estimated, and may be larger or smaller than the long-run average. The discrepancy between the option value and the long run average results in a PE specifically related to stimulus presentation (Fig. S3, Left Frame). Second, due to the time pressure manipulation in the task, responses are generated in a noisy, probabilistic fashion: even though a subject may *intend* to select one option, it may be the case that the other option is selected due to noise in the response process. On trials in which the difference between two options is low, evidence in favor of each option is about the same, and thus the selection of one option over the other entails PE regardless which option is picked. On the other hand, when value differences are extreme, evidence overwhelmingly favors one option over the other, and thus PE is minimal when that option is selected. Together, both stimulus- and response-related (Fig. S3, Right Frame) PEs contribute to the ‘W’ pattern predicted by the PRO model.

**Figure S3.**
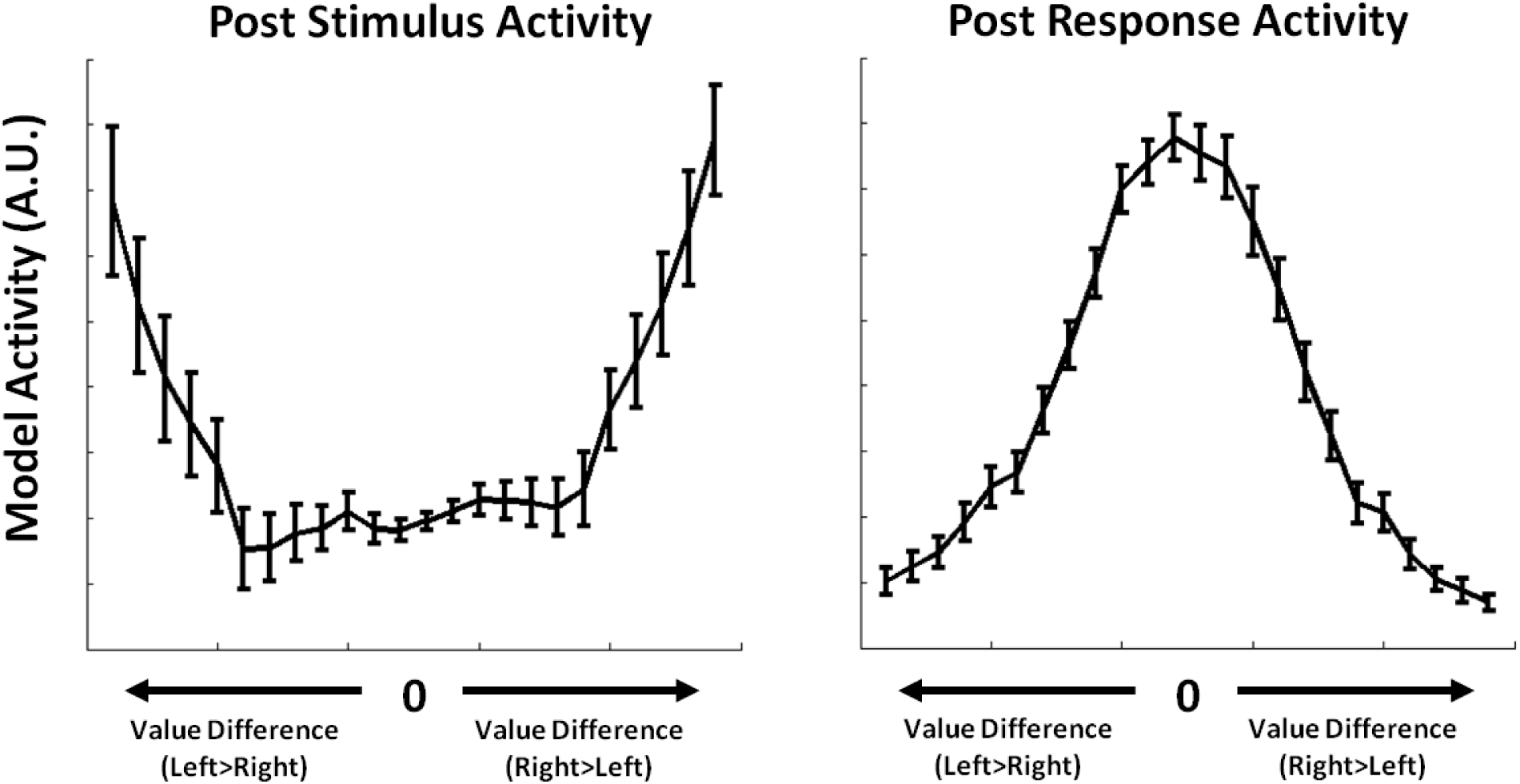
PRO Model Simulated Activity - Following presentation of a stimulus, model activity increases as a function of value differences due to the surprisingly low value of one option. Following a response, PRO model activity increases with the similarity of option values - choosing either response entails increased surprise at *not* choosing the alternative response.

Finally, PEs are generated following task-related feedback. Once an option has been selected, subjects have a 50% chance of receiving the points associated with either of the fractal images that made up that option. The discrepancy between the points actually received and the possible points results in a feedback-related PE.

### Alternative formulation of the PRO model

Like the EVC model, the PRO model is based on standard reinforcement learning models; in particular, the PRO model uses temporal difference (TD) formulations^7^ in order to capture temporal dynamics of activity in ACC^5^. While the use of TD learning allows the model to capture a range of effects in ACC related to temporal contingencies^8^ and learning^9^, it also involves additional model complexity that may obscure a direct comparison with the EVC model. In order to facilitate such a comparison, and to demonstrate that the predictions of the PRO model do not depend on the additional complexity involved in learning and tracking temporal contingencies, we provide a simplified formulation of the PRO model using equations substantially similar to those used to specify the EVC model.

The two core functions of the PRO model can be distilled to prediction and prediction error (PE)^5,10^, and these functions apply generally to salient sensory events^9,11^. In our speeded decision-making task, there are three distinct points in each trial in which such a salient event occurs: 1) the delivery of task feedback in the form of points awarded for a trial, 2) the generation of a response, and 3) the presentation of task stimuli. The range of effects attributed to dACC by the PRO model are described by “negative surprise” – the negative component of PE^5^ – following the occurrence of a salient event:

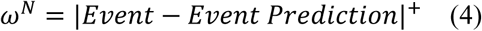

where *ω*^*N*^ is the negative component of the PE associated with the non-occurrence of a predicted event. The functional form of eq. 4 remains identical for all salient, predictable events; in order to derive predictions from the PRO model, then, it is only necessary to specify what is being predicted at each point in the speeded decision-making task. At the delivery of task feedback, the model predicts the amount of points to be received based on the selected option. As noted above, the predictions of the model may be biased by the salience/value of possible outcomes – model predictions are therefore proportional to the expected value:

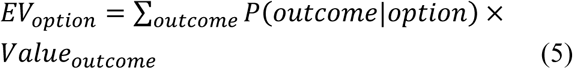

Response-generation in our simplified version of the PRO model is modeled using a sigmoid function, similar to that used in our implementation of the EVC model

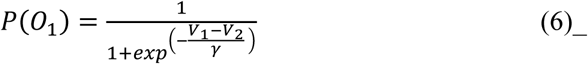

with the exception that there is no effort term and β is subsumed by the γ term. As with feedback, choices are informed by the salience/value of that choice:

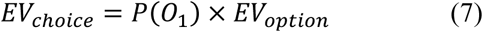

Finally, over the entire task, each option will have a long-run average value of around 45 points. That is, given all possible combinations of stimuli, the long-term expected value of an option before observing the stimuli themselves will be 45 points. Prior to the presentation of task stimuli, therefore, the model predicts that, on average, this average value:

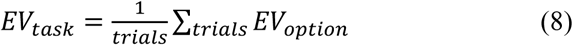

The quantities in eqs 5, 7 & 8 are compared with the actual event that occurred at each salient point in each trial as in eq. 4 in order to generate model predictions. Model activity is therefore the product of only a single equation that applies over a broad range of possible inputs.

